# CAP peptide artificially induces insect gall

**DOI:** 10.1101/2024.01.06.574462

**Authors:** Tomoko Hirano, Tomoaki Sakamoto, Seisuke Kimura, Takumi Nakayama, Mitsuhiko P. Sato, Kenta Shirasawa, Masa H. Sato

## Abstract

Galls caused by gall-inducing insects in their host plants clearly illustrate the concept of ‘extended phenotype’, which refers to traits expressed in a host organism when manipulated by a parasite. Candidate effector molecules involved in gall formation, such as phytohormones, amino acids, and proteins, have been reported in numerous studies. However, to date, no attempts to artificially regenerate gall structures using effector candidates have been reported. In this study, we tested the peptide from <u>C</u>ysteine-rich secretory proteins, <u>A</u>ntigen 5, and <u>P</u>athogenesis-related 1 proteins, CAP peptide as a gall-inducing effector candidate obtained from transcripts isolated from the horned gall aphid, (*Schlechtendalia chinensis*) through in silico screening and the Arabidopsis-based gall-forming assay, which is a bioassay system for analysing the molecular mechanisms of gall formation. Furthermore, we succeeded in generating an artificial gall in the host plant *Veronica peregrina,* without any insect parasitism, using three minimal effector elements: CAP peptide, auxin, and cytokinin. Given the strong similarities observed in organ structure with a central cavity and three types of tissue and gene expression patterns between the native and artificial galls, we concluded that CAP peptide is a general gall-inducing effector peptide secreted by gall-inducing insects.

## Main text

Insect galls induced by gall-inducing insects, gall midges, aphids, and cynipids in their host plants for shelter and nutrients are unique, complex, and highly organised structures; (i) callus-like cells transformed from palisade and spongy tissues used for insect feeding in inner-layer cells (ii) protective outer tissue composed of lignified secondary cell walls for shelter and protection in outer-layer cells, and (iii) vascular tissues which transports water and nutrients to the callus-like cells in the gall ^1,2^ . Gall-inducing insects reportedly control the host developmental programs by injecting certain effectors, including phytohormones, into host cells to produce galls.^3,4^ Recent transcriptome analysis of several insect galls has revealed that gall-inducing insects manipulate plant floral organ-developmental programs to convert cells in source tissues into flower- and fruit-like sink tissues by triggering the initiation of reproductive development.^5–8^

Effector molecules involved in gall induction, including phytohormones such as auxins and cytokinins, amino acids, and proteins, have been extensively researched in the salivary secretions and oviposition fluids of larvae, parental insects, or both.^9–12^ However, actual effector molecules and the molecular mechanisms underlying insect gall formation remain largely unknown. In this study, we identified a novel gall-inducing effector candidate, the CAP peptide, from the gall-inducing aphid *Schlechtendalia chinensis*, commonly known as the horned gall aphid. Furthermore, by applying the highly conserved six-amino acid region (F/Y-T-Q-I/V-V-W) of the CAP protein in combination with phytohormones, we artificially generated a gall structure in *Veronica peregrina,* which is a host plant of the gall-inducing weevil *Gymnaetron miyoshii* Miyoshi.

Using the <u>A</u>rabidopsis-<u>b</u>ased <u>gal</u>l-forming <u>a</u>ssay (Ab-GALFA), we previously showed that the extract of the gall-inducing aphid, *S. chinensis*, induced gall-like structures in the root tip region of Arabidopsis,^13^ suggesting that the insect extract may contain unknown gall-inducing effector molecules.

To identify candidate effector molecules, we first performed Ab-GALFA to determine the basic characteristics of these potential effector molecules. Arabidopsis seedlings were incubated for 16 h in an insect-extracts pre-treated at 99 °C for 10 min or digested with Actinase E. Gall-inducing activity was shown only by the heat-treated solution containing aphid extracts, suggesting that the effector would be a small protein or a peptide (Extended Data Fig. 1). *S. chinensis* induces gall formation in the host plant through the following process: after a single *S. chinensis* fundatrix initiates gall formation in the wing leaf of host plant *Rhus javania,* pale-yellowish apterous viviparae clones continuously increase their population inside the growing galls from May to September. Subsequently, in late September, the body of the viviparae becomes brownish in colour, and the galls stop growing.^14^ Based on the gall growing process and the colouration of the aphid bodies, we performed RNA-seq analysis of the three stages in *S. chinensis*, fundatrix, pale-yellowish apterous viviparae, and brownish ones, and set the following *in silico* screening criteria for identifying potential gall-inducing factors isolated from *S. chinensis*: (1) the upregulated genes (sum >10, log FC >3, and false discovery rate [FDR] <0.05) in both fundatrixs and pale-yellowish apterous viviparae, compared with those in brownish ones; (2) the transcripts were subjected to BLAST (Basic Local Alignment Search Tool) search in the Arabidopsis genome database (TAIR10) to identify proteins similar to those found in Arabidopsis; and (3) candidate translations having an N-terminal signal peptide were identified using SignalP (SignalP 3.0: https://services.healthtech.dtu.dk/service.php?SignalP-3.0).

In the first step, 3,986 genes were identified among 132,421 *S. chinensis* differentially expressed genes (DEGs), among which, 1,454 genes complied with criterion (2), and 497 genes were identified using criterion (3). Furthermore, among these 497 targets, 244 belonged to the same family: cysteine-rich secretory proteins, antigen 5, and pathogenesis-related 1 (CAP) proteins (Fig. 1a).

**Fig. 1.**
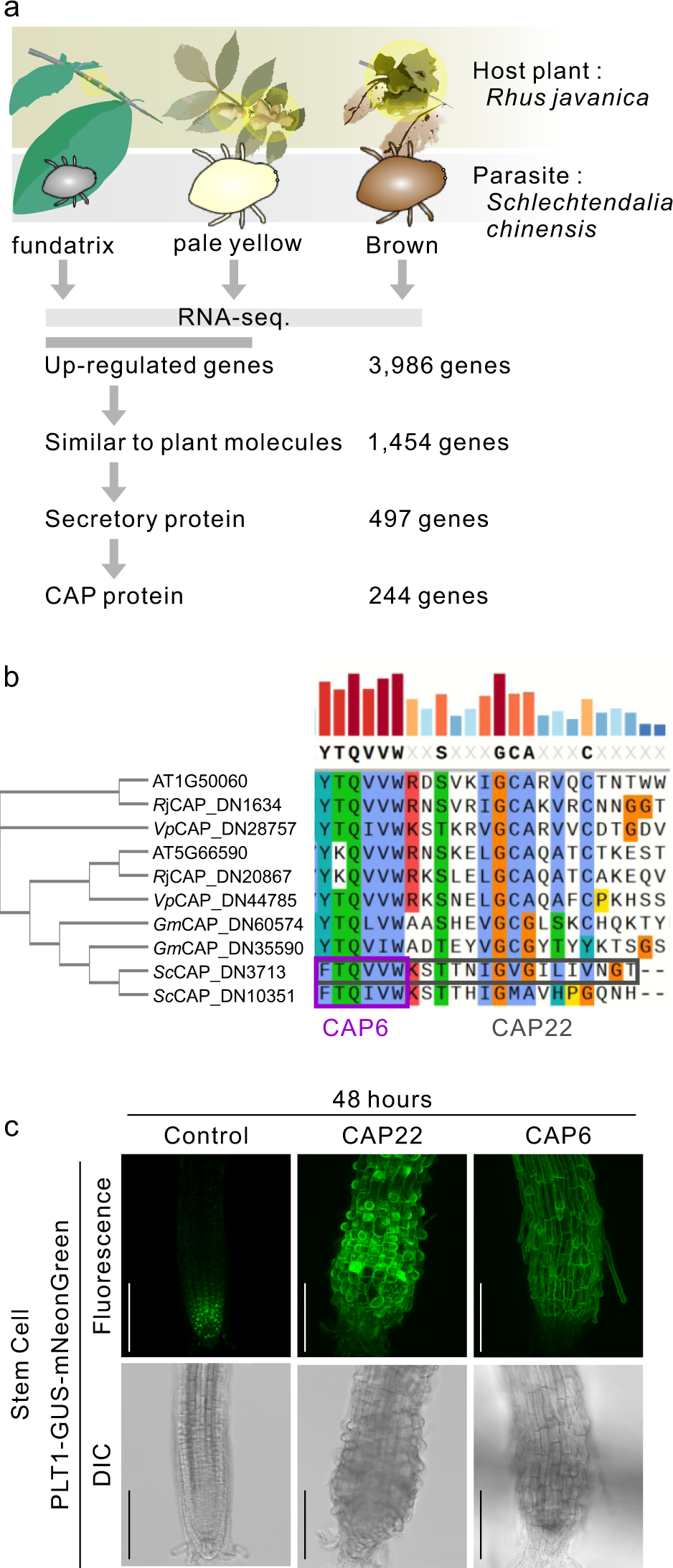
**(a)** Scheme of gall formation-effector screening. Among the upregulated genes in fundatrix and pale-yellow larva compared to brown larva, those with homologues in Arabidopsis were selected. Subsequently, genes encoding proteins with secreted signals were selected, and the candidate was CAP gene/protein. **(b)** Phylogenetic tree and partial alignment of the CAP protein sequences of *R. javanica* (*Rj*)*, S. chinensis* (*Sc*), Arabidopsis thaliana (AGI code), *Veronica peregrina* (*Vp*), and *Gymnetron miyoshi* (*Gm*) revealed a highly conserved 6-amino acid sequence, named CAP-p6, and a 22-amino acid sequence consisting of CAP-p6 core sequence plus additional 16 amino acids, named CAP-p22. **(c)** Images of mNeonGreen (green) or DIC (grey) in PLT1-GUS-mNeonGreen seedlings with or without treatment of 4 µM CAP6 or CAP22 peptides for 48 h are represented as projection images of serial z-stack sections. Bars = 100 μm.

The CAP superfamily is conserved across a broad range of eukaryotes, including insects and higher plants.^15^ Furthermore, CAP superfamily proteins are defined by the presence of four highly conserved motifs designated CAP1–CAP4.^16^ In this study, we specifically aligned CAP family proteins from two pairs of the parasitic insect and the host plant, *S. chinensis* and *Rhus javania*, *Gymnaetron miyoshii* Miyoshi and *V. peregrina*, and the model plant *Arabidopsis thaliana*, and found that the CAP1 core sequence (F/Y-T-Q-I/V-V-W) near the C-terminal region is highly conserved among insects and plants (Fig. 1b and Extended Data Fig. 2). Hence, we speculated that the conserved region might possess certain biological activities affecting higher plants; therefore, we chemically synthesised the 6 (CAP-p6) and 22 (CAP-p22) polypeptides containing the conserved core sequence to test for any biological activity of these polypeptides using Ab-GALFA (Fig. 1b). First, we investigated whether both peptides, CAP-p22 and CAP-p6, have the same ability to induce gall-like structures in Arabidopsis roots. This was done using a stem cell marker-line expressing PLETHORA1 (PLT1) fused with mNeonGreen under the control of the PLT1 promoter (PLT1p:PLT1-GUS-mNeonGreen), because PLT1 was shown to be ectopically expressed during the formation of a gall-like structure.^13^ Treatment with 4 μM CAP-p22 or CAP-p6 peptides for 48 h induced PLT1-mNeonGreen fluorescence in the root tip region including, the division, transition, and elongation zones, indicating that both CAP-p22 and CAP-p6 have the same ability to induce stem cell characteristics in a broad area of the root tip region (Fig. 1c).

Subsequently, we performed RNA sequencing (RNA-seq) in 4-day-old Arabidopsis seedlings subjected to four different treatments (4 μM CAP-p22 peptide for 4, 16, and 48 h and 60 nM CAP-p22 peptide for 16 h) for a comprehensive analysis of gene expression. We identified DEGs (sum >10, |log FC| >1, and FDR <0.05) at each sampling time-point relative to time 0. The results showed a total of 4,477 DEGs (1,315 upregulated and 3,162 downregulated), 793 DEGs (128 upregulated and 665 downregulated), and 1,695 DEGs (809 upregulated and 886 downregulated), induced by 4 μM CAP-p22 peptide treatment in Arabidopsis seedlings treated for 4, 16, and 48 h, respectively. These DEGs were categorised into ‘gene expression’, ‘development’, ‘respiration’, and ‘photosynthesis’ on Gene Ontology (GO) term enrichment analysis. In contrast, 4,337 DEGs (678 upregulated and 3,659 downregulated) induced by treatment with a low CAP peptide concentration (60 nM) for 16 h were categorised into ‘photosynthesis’, ‘abiotic stress tolerance’, ‘defense’, and ‘circadian rhythm’(Fig. 2a). Thus, peptide CAP-p22 induced different gene expression patterns in a dose-dependent manner in Arabidopsis seedlings.

**Fig. 2.**
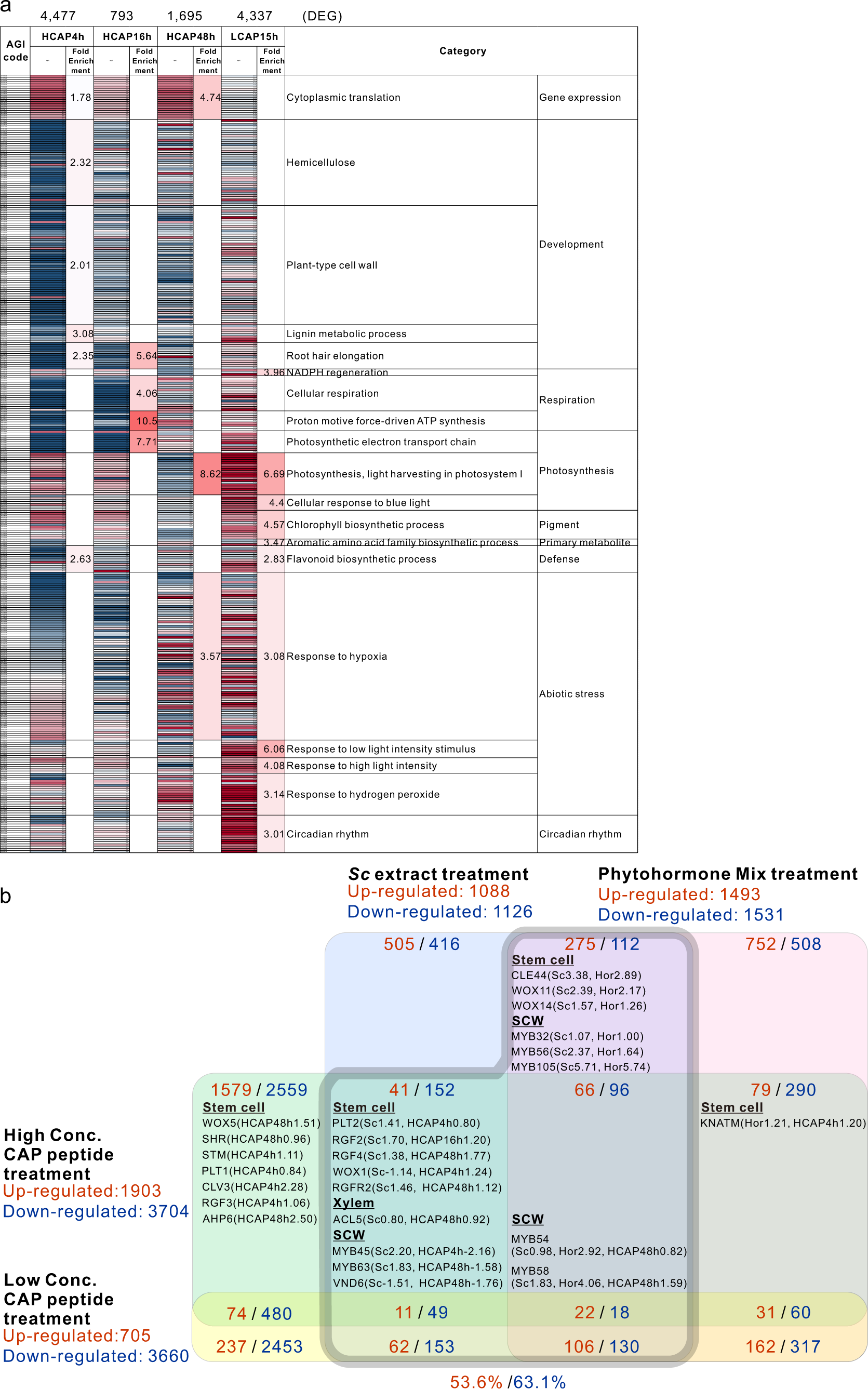
**(a)** Upregulated/downregulated genes (sum>1, |log FC|>1, and FDR<0.05) in Arabidopsis seedlings with 4 µM or 60 nM CAP-p22, compared to without peptide were analysed using gene ontology (GO) term enrichment analysis. **(b)** Venn diagram showing the overlap of upregulated/downregulated genes between the treatment groups of CAP peptide of high (4 µM) or low (60 nM) concentration (sum>1, |log FC|>1, and FDR<0.05), Aphid extract (sum>10, |log FC|>1, FDR<0.05), Phytohormone mixture (sum>10, |log FC|>1, FDR<0.05), showing the marker genes associated with the stem cell, the secondary cell wall (SCW), and the xylem.

The genes involved in ‘stem cell’, ‘secondary cell wall’, and ‘xylem formation’ are reportedly induced during the initial stage of the gall formation process.^6–7^ Furthermore, the same genes were expressed during the formation of the gall-like structure in Arabidopsis seedlings treated with the gall-inducing aphid extract in combination with phytohormones but not when treated with phytohormones in combination with aphid bodies. This suggests that gall formation requires not only phytohormones but also an unidentified gall-inducing factor.^13^ As treatment with 4 μM peptide CAP-p22 effectively induced the expression of stem cell-formation genes (e.g. *WUSCHEL RELATED HOMEOBOX [WOX] 5*, *SHOOT ROOT [SHR]*, *SHOOT MERISTEMLESS [STM]*, *PLETHORA [PLT] 1*, *CLAVATA-RELATED [CLE] 3*, *ROOT MERISTEM GROWTH FACTOR [RGF] 3*, *HISTIDINE PHOSPHOTRANSFER PROTEIN [HP] 6*, *PLT 2*, *RGF 2*, *RGF 4*, and *WOX* 1), secondary cell wall marker genes (e.g. *MYB45*, *MYB63*, *MYB54*, and *MYB58*), and xylem marker genes (e.g. *ACAULIS [ACL] 5*), we inferred that CAP-p22 treatment partially induced the gene expression pattern responsible for gall formation (Fig. 2b). The total number of genes induced by CAP or the artificial phytohormone mixture treatment was over 53 % of all the genes induced by gall-inducing aphid extracts (Fig. 2b). Therefore, we investigated whether the CAP peptide has any gall-inducing effector activity using Ab-GALFA. We previously measured the contents of phytohormone, indole acetic acid (IAA), isopentyladenine (iP), and trans-zeatin (tZ), abscisic acid (ABA), salicylic acid (SA), jasmonic acid (JA), jasmonoyl-L-isoleucine (JA-Ile) in *S. chinensis* and reproduced an artificial phytohormone mixture (AHM) that replicates this composition.^13^ A gall-like structure was induced at the root tip through treatment with a mixture of CAP-p6 plus AHM and CAP-p6 plus indole-3-acetic acid (IAA) and 6-benzylaminopurine(BAP) (at the same concentration as in AHM), as well as the aphid extract after treatment for 16 h (Fig. 3a, Extended Data Fig. 3a and 3b). Under these conditions, fluorescence of the stem cell-marker gene PLT1-mNeonGreen was observed not only in the division zone but also in the elongation zone of the root epidermal cells, indicating that the stem cell niche spread from the meristematic region to the elongation zone in the root tip (Fig. 3a). Conversely, these structures were not induced by treatment with AHM, IAA plus BAP, or CAP-p6 peptide alone (Fig. 3a). Consequently, the secondary cell wall emerged in the epidermal cells of the root transition and elongation zones (Extended Data Fig. 3a and 3b), and the xylem vessel extended to the root tip region after treatment with IAA and BAP, as well as the aphid extract after treatment for 16 h (Extended Data Fig. 3a and 3c). These results indicate that peptide CAP-p6, auxin, and cytokinin are the minimum components required for successful gall-like structure formation in the root tip region of Arabidopsis, clearly implying that the CAP peptide acts as a gall-inducing effector in plants.

**Fig. 3.**
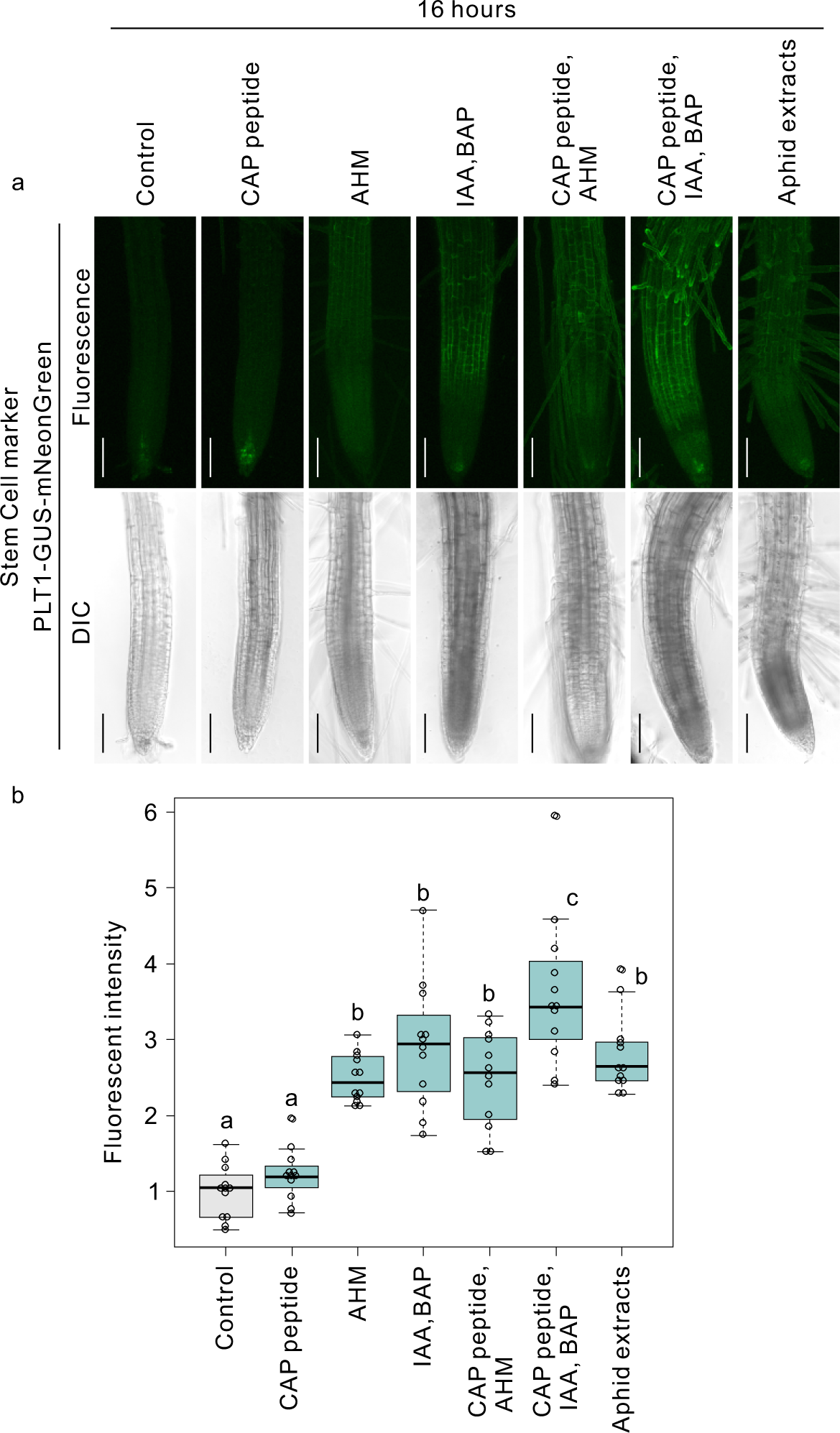
Localisation of PLT1-GUS-mNeonGreen in 4-day-old seedlings of the PLT1-GUS-mNeonGreen line treated with or without 5 μM CAP-p6 peptide, artificial hormone mixture (AHM), a mixture of IAA and BAP, a mixture of CAP peptide and AHM, and a mixture of and CAP peptide, IAA, and BAP, and Aphid extract for 16 h (**a**) and measured fluorescence intensity **(b)**. Groups that have different letters are significantly different from each other (n = 12 each, P < 0.05, Wilcoxon and Steel–Dwass tests). Bars = 100 μm.

Subsequently, we attempted to artificially reproduce a real gall structure without the intervention of insect parasitism in *V. peregrina,* an annual plant that forms small bud-like galls at the shoot meristem upon attack by the gall-inducing weebill, *Gymnaetron miyoshii Miyoshi* (Extended Data Fig. 4a,b, Fig. 4a). *V. perigrina* seedlings were cultured on 1/2 MS medium containing 0, 5, or 10 μM of CAP-p6 peptide; 0.05, 0.1, or 0.2 mg/mL of 1-naphthaleneacetic acid (NAA); and 0.05 or 0.25 mg/mL of 6-benzylaminopurine (BAP). The results showed that, after four weeks, a bulb-shaped structure (Fig. 4b, Extended Data Fig. 4) resembling the native *G. miyoshii Miyoshi-*induced gall at the stem tips, was formed at the shoot apex of *V. perigrina* (Fig. 4a). In contrast, plants cultured on medium containing phytohormones alone formed roots and leaves after callus formation, while those cultured in medium containing CAP peptide alone grew normally (Extended Data Fig. 5). Subsequently, we compared the structure of a native gall (Fig. 4a,b) with this bulb-shaped structure (Fig. 4c,d) formed at the shoot apex of *V. perigrina* by preparing paraffin sections. We found that they shared four common characteristics of the gall structure: (i) the central cavity for the living space of the insect, (ii) the presence of callus-like or parenchyma cells in the inner layer of the gall, (iii) the secondary cell wall formed in the cells of the outer layer (Fig. 4c,d), and (iv) tracheary elements emerged among the callus tissue cells (Fig. 4e,f).

**Fig. 4.**
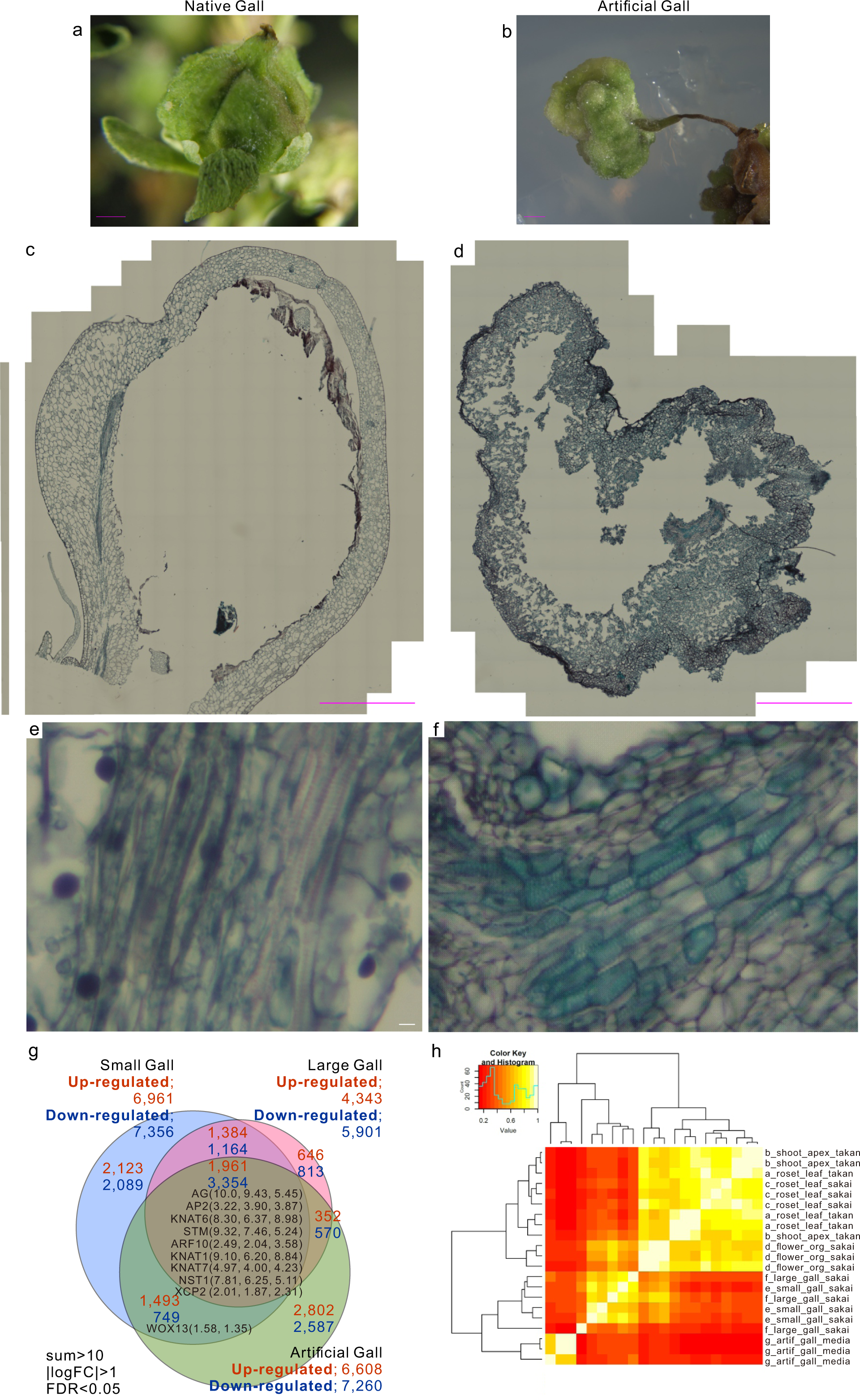
A gall of *Veronica peregrina* growing in Sakai-fureainomori, Osaka, Japan **(a)** and the artificial gall of one **(b)** were historically stained with Safranin/O, Fast Green, and haematoxylin **(c, d, e, f)** with tracheary elements (TEs) in the xylem **(e, f)**. Venn diagram showing the overlap of upregulated and downregulated genes **(g)** (sum>10, |log FC|>1, and FDR<0.05) between native small galls, large galls, and artificial galls, showing the log FC values of genes commonly expressed in the insect gall of different plants and the gall-like structure of Arabidopsis by Ab-GALFA (**g**). (**h**) Hierarchical clustering of a gene–gene expression correlation matrix from different tissues: the rosette leaf (a); shoot apex (b) in *Veronica peregrina* from Takano; and the rosette leaf (c), the flower (d), the native small gall (e), the native large gall (f), and the artificial gall (g) in *Veronica peregrina* from Sakai. Bars = 1 mm (**a–d**) and 10 μm (**e** and **f**).

Furthermore, to investigate the similarity of gene expression patterns between the native gall and the bulb-shaped structure, we performed genome sequencing and RNA-seq analysis of *V. perigrina* to compare the expression profiles of the native and bulb-like structures artificially formed by the gall-inducing elements, CAP peptide, auxin, and cytokinin.

Briefly, 34.2 Gb HiFi reads of *V. perigrina* genomic DNA, totalling 38.0× the estimated genome size (899.1 Mb), were assembled into 143 contigs spanning 822.7 Mb with an N50 of 24.3 Mb (Extended Table 1). The complete Benchmarking Universal Single-Copy Orthologs (BUSCO) score, indicating completeness of the assembly, was 98.8 % (Extended Table 2), consisting of 18.8 % and 80.0 % single-copy and duplicated scores, respectively. RNA-Seq analysis was performed on young leaves, native small galls (3–5 mm in diameter), native large galls (5–10 mm in diameter), and the bulb-like structures of *V. perigrina*. The RNA-seq reads were mapped to the *V. perigrina* genome to predict 62,545 high-confidence protein-coding genes (Extended Table 1), which included 96.6 % complete BUSCO scores (Table 2). Repetitive sequences comprised 54.3 % of the genome (Extended Table 3).

We used RNA sequencing data of small and large native galls and artificial galls to examine the similarity of the gene expression patterns to set the selection criteria of DEGs at an FDR value <0.05 and |log FC|> 1. In total, 14,317, 10,244 and 13,868 DEGs were identified in the native small, native large, and artificial galls, respectively, including 6,961, 4,343, and 6,608 upregulated and 7,356, 5,901, and 7,260 downregulated genes, respectively.

In total, 3,806 DEGs from the 6,608 upregulated genes and 4,673 DEGs from the 7,260 downregulated genes in the artificial gall overlapped with the corresponding DEGs in the native small or large galls (Fig. 4g). These genes were 57.6 % and 64.3 % of all upregulated and downregulated genes expressed in artificial galls, respectively, and were included among common expression genes extracted from transcriptome data of developing galls of *A. montana, G. obovatum, R. javanica*, and *V. riparia*,^5–7^ the genes associated with stem cell generation and floral organ development, *WOX* family, *KNOX* family, *STM*, *AGAMOUS* (*AG*), and *APETALA* (*AP*) 2. Furthermore the artificial and native galls exhibited a strong similarity in the hierarchical clustering of a gene–gene expression correlation matrix in RNA-seq data of different tissues: the rosette leaf; shoot apex in *V. peregrina* from Kami-Takano in Kyoto City; and rosette leaf, flower, native small gall, native large gall, and artificial gall in *V. peregrina* from Sakai (Fig. 4h).

Gall effector candidate molecules were selected based on the data of transcriptome and phytohormone analyses and Ab-GALFA using the gall-inducing insect, *S. chinensis*. The three elements combined, CAP peptide, auxin, and cytokinin, induced an artificial gall with a gene expression patter that shared more than 57.6 % identity with the native gall in gall-forming *V. peregrina*. The bulb-like structure induced at the shoot apex by the CAP peptide and phytohormone combination treatment was an artificial gall with strong similarities in organ structure and gene expression patterns to the native gall. Based on these results, we concluded that CAP is a general gall-inducing effector secreted by gall-inducing insects.

Therefore, we succeeded in artificially generating a gall structure on the stem apex of *V. perigrina* by adding CAP peptide, auxin, and cytokinin to 1/2 MS medium in the absence of any insect parasitism. Further studies are needed to identify native secreted CAP peptides from gall-inducing insects and to demonstrate that CAP peptides can function as general gall-inducing effectors in various gall-forming host plants.

## Materials and Methods

### Plant material and growth conditions

*Arabidopsis thaliana* ecotype Columbia (Col-0) was used as the wild type (WT) in all experiments. Arabidopsis seeds were surface sterilised and germinated on 1/2 Murashige–Skoog (MS) 1.2 % agarose plates, and the plants were grown on tilted plates at 45°. Arabidopsis plants were grown under white light with a 16/8 h (light/dark) photoperiod at 22 °C for 4 days before Ab-GALFA and confocal microscopy observation. Transgenic marker lines in the PLT1pro::PLT1-GUS-mNeonGreen plasmid expressing PLT1, β-glucuronidase (GUS) reporter, and Bright Fluorescent Protein mNeonGreen under 3000 bp own promoter region were generated. *V. peregrina* samples for WGS and RNA-seq were collected from Sakai-fureainomori natural park, Osaka, Japan (Japan; 34°27’30.2’’N; 135°31’ 29.9’’E).

### Arabidopsis-based gall formation assay (Ab-GALFA)

Ab-GALFA was performed as described by Hirano et al.^13^ Approximately 10 *S. chinensis* bodies isolated from the rapid growth phase of *R. javanica* gall, and 10 bodies of non-galling aphids *Acythosiphon pisum* and *Megoura crassicauda* were frozen in liquid nitrogen, ground to a fine powder, and suspended in 50 µL of sterile distilled water (DW), then the suspensions were centrifuged at 20,000xg for 5 min at 4°C to prepare the *S. chinensis* (Sc) extract stock solution. The working solution of aphid extracts was prepared by diluting the stock solution 10-fold. Arabidopsis seedlings grown vertically on 1/2 MS medium with 1.2% agarose for 4 days were gently removed from the agarose surface using micropipette tips, placed in a Petri dish, and the entire bodies of seedlings were incubated with different concentrations of Sc, Ap or Mc extracts or sterilized water (a negative control) under normal growth conditions for ≥ 15 h. Following incubation, the seedlings were used for subsequent analyses.

### Histological staining of seedlings

To stain the components of the secondary cell wall, Arabidopsis seedlings were washed in sterilised water and incubated in 1 μg/mL WGA-Alexa Fluor 488 (Invitrogen) in sterilised water at 25 °C for 1 h and washed five times for 30 min. Lignin and suberin were stained with basic fuchsin and Auramine O, respectively, as described by Ursche et al.^17^

### Histological staining of Veronica galls

For native and artificial galls, 10 mm^3^ samples were fixed in the formalin-acetic-70 % alcohol (FAA, v/v) buffer and exhausted with an aspirator pump. Serial transverse and longitudinal sections of paraffin-embedded tissues were sequentially stained with safranin, fast green, and haematoxylin. Subsequently, the sections were observed under an AXIO Zoom.V16 (Zeiss).

### Image analysis

Fluorescence and differential interference contrast (DIC) images were obtained using a Leica TCS SP8 laser scanning confocal microscope. Captured images were processed using Leica LAS X or ImageJ software. For quantitative analysis, all image data in each experiment were obtained at an identical laser power and detector sensitivity. All experiments were repeated at least thrice, and the same results were obtained. Analysed image data are shown as box-and-whisker plots. Boxes and solid lines in the box-and-whisker plots show the upper (75^th^) and lower (25^th^) quartiles and median values, respectively. Whiskers indicate 95 % confidence intervals. Significant differences were evaluated using the Mann–Whitney U test.

### Statistical analysis

Boxes and solid lines in the box-and-whisker plots show the upper (75^th^) and lower (25^th^) quartiles and median values, respectively. Whiskers indicate the 95 % confidence intervals. Significant differences (P<0.05) between samples were evaluated using the Wilcoxon and Steel–Dwass tests .

### Measurements of intensity and statistical analysis

Subsets of data with defined x-, y-, and z-dimensions were acquired using LASX software (Leica, Germany), and all subsets were transformed into 2D images using the maximum intensity projection function of the LASX software. Uniform brightness and contrast corrections were performed on all images before they were exported for analysis. All quantitative data were produced using the publicly available software, Image Studio (LI-COR). Values were normalised to the average fluorescence intensity of the control group. Final statistical data evaluation and plot preparation were performed using the R software.

### RNA extraction, library construction, and RNA sequencing

Total RNA was extracted from the plant tissues using the RNeasy Plant Mini Kit and Qiagen RNase-Free DNase Set (Qiagen, Hilden, Germany). Three independent RNA samples from each tissue were used for analysis. The RNA quality was assessed by determining the RNA integrity number using an RNA 6000 bioanalyzer and an RNA Nano Chip (Agilent Technologies). RNA-seq libraries were prepared using the Illumina TruSeq® Stranded RNA LT kit (Illumina, San Diego, CA, USA) according to manufacturer’s instructions. Pooled libraries were sequenced using NextSeq 500 (Illumina), and single-end reads of 75 bp length were obtained. Reads were mapped to the reference using TopHat2.^18^ The HTSeq-count script in the HTSeq library was used to count the reads.^19^ Count data were subjected to a trimmed mean of M-value normalisation using EdgeR.^20,21^ DEGs were defined using the EdgeR GLM approach,^20^ and genes with FDRs <0.05 were classified as DEGs.

### Whole-genome sequencing and assembly

Fresh leaves were collected from a single plant of *V. peregrina* and ground under liquid nitrogen to extract genomic DNA using Qiagen Genomic-Tip (Qiagen, Hilden, Germany). A short-read sequence library was prepared using the PCR-free Swift 2S Turbo Flexible DNA Library Kit (Swift Sciences, Ann Arbor, MI, USA) and sequenced on a DNBSEQ G400 (MGI Tech) in paired-end, 150 bp mode. Genome size was estimated using jellyfish (k = 21).

Simultaneously, a long-read sequence library was prepared using the SMRTbell Express Template Prep Kit 2.0 (PacBio, Menlo Park, CA, USA) and sequenced using a PacBio Sequel II system (PacBio, Menlo Park, CA, USA). The obtained HiFi reads were assembled using Hifiasms with default parameters. Assembly completeness was evaluated by Benchmarking Universal Single-Copy Orthologs (BUSCO) using the lineage dataset embryophyta_odb10.

### Gene prediction and repeat sequence analysis

To annotate protein-coding genes, we combined RNA-based, homology-based, and ab initio methods. The RNA-seq reads were mapped onto the genome assembly using HiSat2,^22^ and the resultant BAM files were used for gene prediction using BRAKER2.^23^ The resultant gene models were functionally annotated using Emapper implemented in EggNOG (a cutoff value of <1E-10) and DIAMOND search^24^ against the UniProtKB database (<1E-20).^25^ In addition, the RNA-seq reads were mapped onto the gene models with salmon^26^ to quantify gene expression as TPM. Gene models annotated with both EggNOG and UniProtKB or with TPM values >1 were selected as high-confidence gene models. Repetitive sequences in the assembly were identified using RepeatMasker (https://www.repeatmasker.org), version 4.1.2; parameters: -poly and -xsmall, using repeat sequences registered in Repbase^27^ and a *de novo* repeat library built with RepeatModeler (https://www.repeatmasker.org), version 2.0.2a, with default parameters.

## Data availability

The datasets generated for this study can be found in the DDBJ as the BioProject Submission IDs; PRJDBxxxxx (https://), PRJDB17190, and Plant Garden (https://plantgarden.jp/). All data and materials appearing in this study are available from the authors upon reasonable request.

## AUTHOR CONTRIBUTIONS

T. Hirano and M.H.S. conceived and designed the study. T. Hirano performed almost all experiments in this study. T. K. performed histological experiments. S.T. and S.K. performed RNA-Seq analysis. T. Hirano, S.T., S.K., M.P.S., K.S., and M.H.S. analyzed the data. T. Hirano and M.H.S. wrote the manuscript, and supervised the project.

## Author information

The authors declare no competing financial interests. Correspondence and requests for materials should be addressed to M.H.S.

## ACKNOWLEDGMENTS

We thank Ms. Kaori Mizuno (Kyoto Prefectural University) for technical help for library construction and RNA-Seq, and Y. Kishida, C. Minami, K. Ozawa, H. Tsuruoka, and A. Watanabe (Kazusa DNA Research Institute) for technical assistance. This work was supported by the Japan Society for the Promotion of Science; a-Grant-in-Aid for Scientific Research (A) (Grant No. 19H00933) to M.H.S., by JST, PRESTO Grant Number JPMJPR20D5, Japan to T. Hirano, by JSPS KAKENHI Grant, grant nos. 21H02513 to S.K. and 22H05172 and 22H05181 to K.S., by the MEXT-Supported Program for the Strategic Research Foundation at Private Universities, grant no S1511023 to S.K.

---

**Extended Data Fig. 1.**
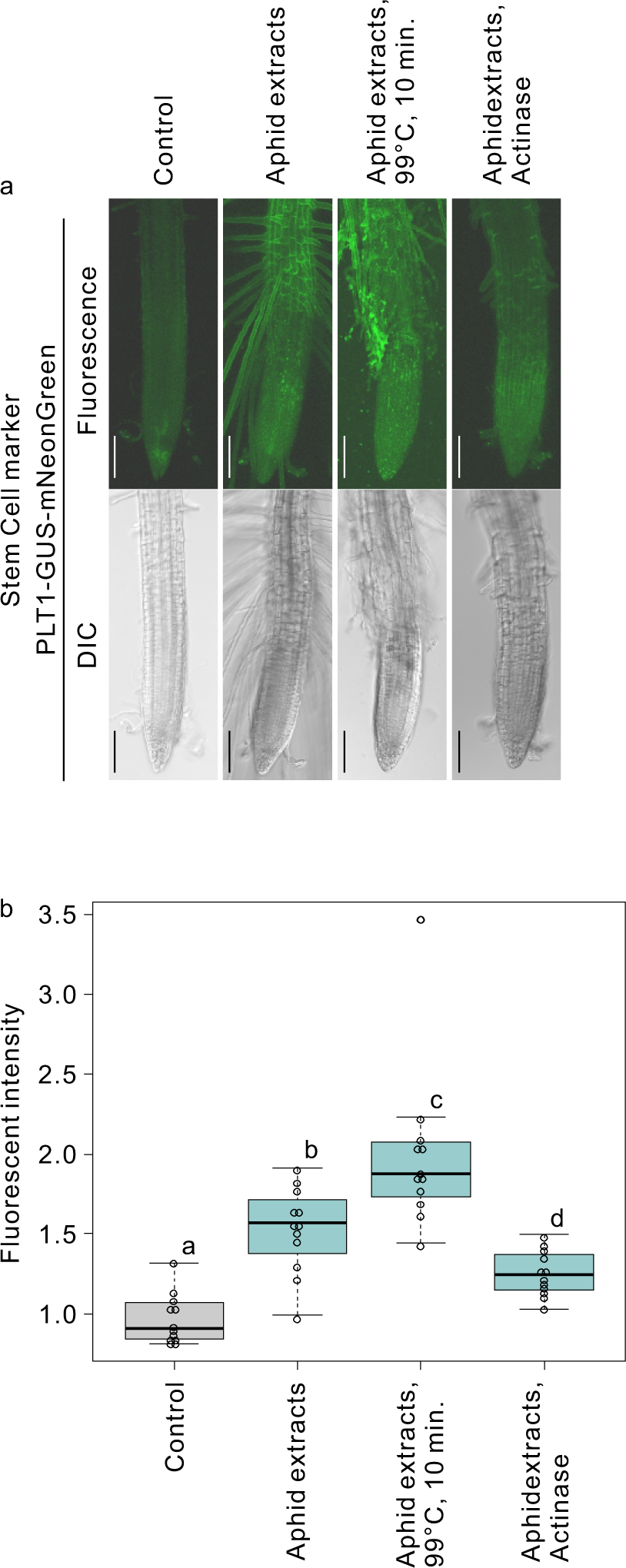
Localisation of PLT1-GUS-mNeonGreen in 4-day-old seedlings of the PLT1-GUS-mNeonGreen line treated for 16 h with or without aphid extract, one boiled with 99 °C for 10 min, and one treated with 1 mg/mL actinase for 5 min (**a**) and measured fluorescence intensity **(b)**. Groups with different letters are significantly different from each other (n = 12 each, P < 0.05, Wilcoxon and Steel–Dwass tests). Bars = 100 μm.

**Extended Data Fig. 2.**
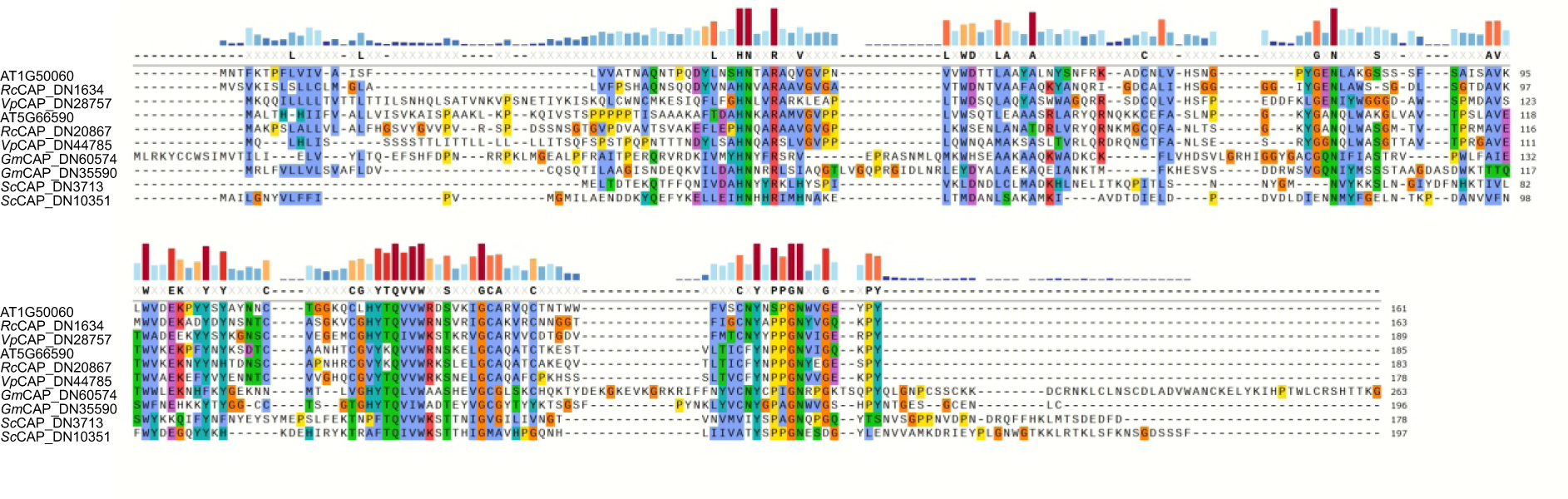
Alignment of the CAP protein sequences of *R. javanica* (*Rj*)*, S. chinensis* (*Sc*), Arabidopsis thaliana (AGI code), *Veronica peregrina* (*Vp*), and *Gymnetron miyoshi* (*Gm*).

**Extended Data Fig. 3.**
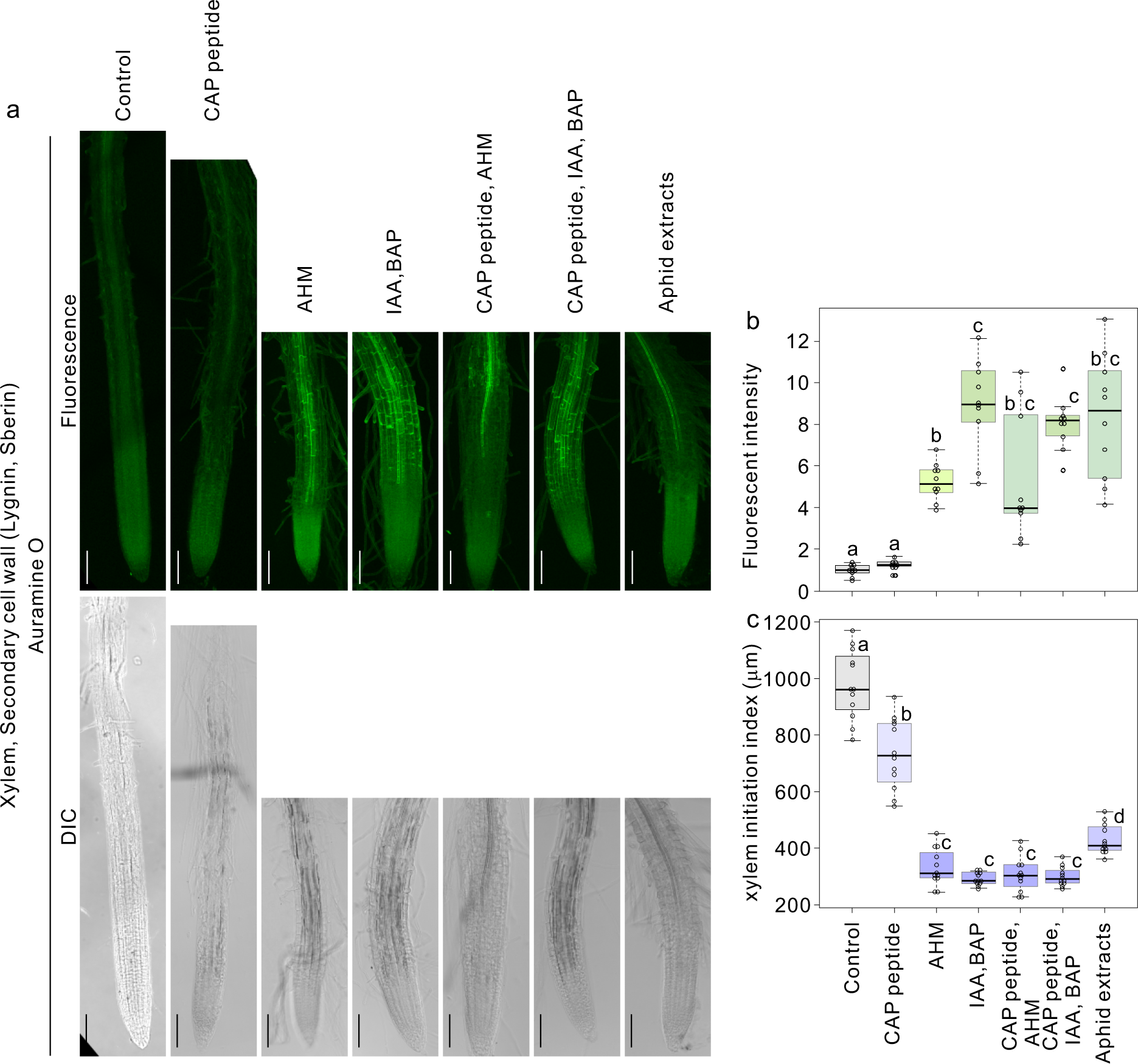
Fluorescent images of auraminO-labelled suberin and lignin in 4-day-old seedlings of Col-0 treated with or without 5 μM CAP-p6 peptide, artificial hormone mixture (AHM), a mixture of IAA and BAP, a mixture of CAP peptide and AHM, and a mixture of and CAP peptide, IAA, and BAP, and Aphid extract for 16 h **(a)** and measured fluorescence intensity **(b)** and the distance between the tip and xylem initiation point **(c)**. Average fluorescence intensity of the control samples was set at 1.0 **(b).** Groups with different letters are significantly different from each other (n = 12 each, P < 0.05, Wilcoxon and Steel–Dwass tests). Bars = 100 μm.

**Extended Data Fig. 4.**
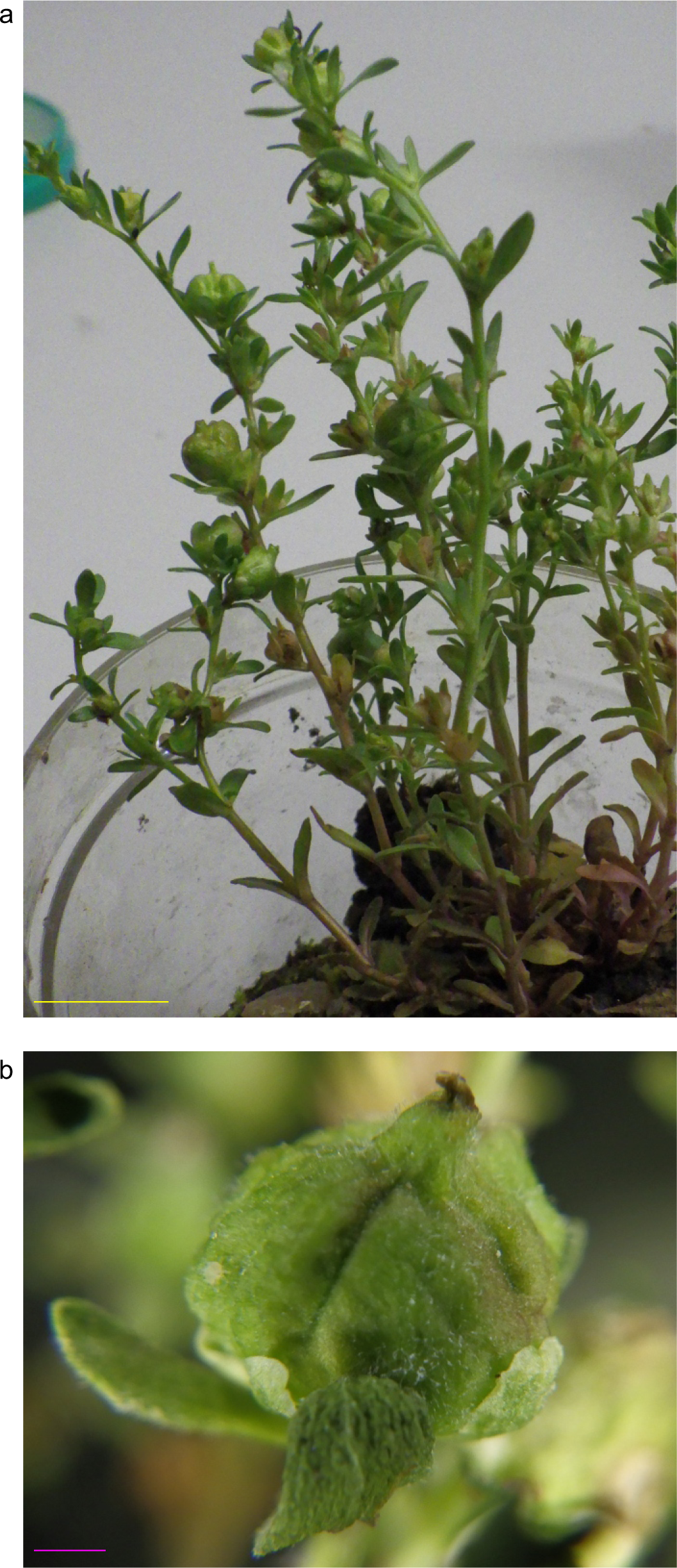
*V. peregrina* growing in Sakai-fureainomori natural park, Osaka, Japan (Japan; 135°31’ 29.9’’E, 34°27’30.2’’N) **(a)** with the gall **(b)** formed in the stem tip. Bars = 10 mm (**a**) and 1 mm (**b**).

**Extended Data Fig. 5.**
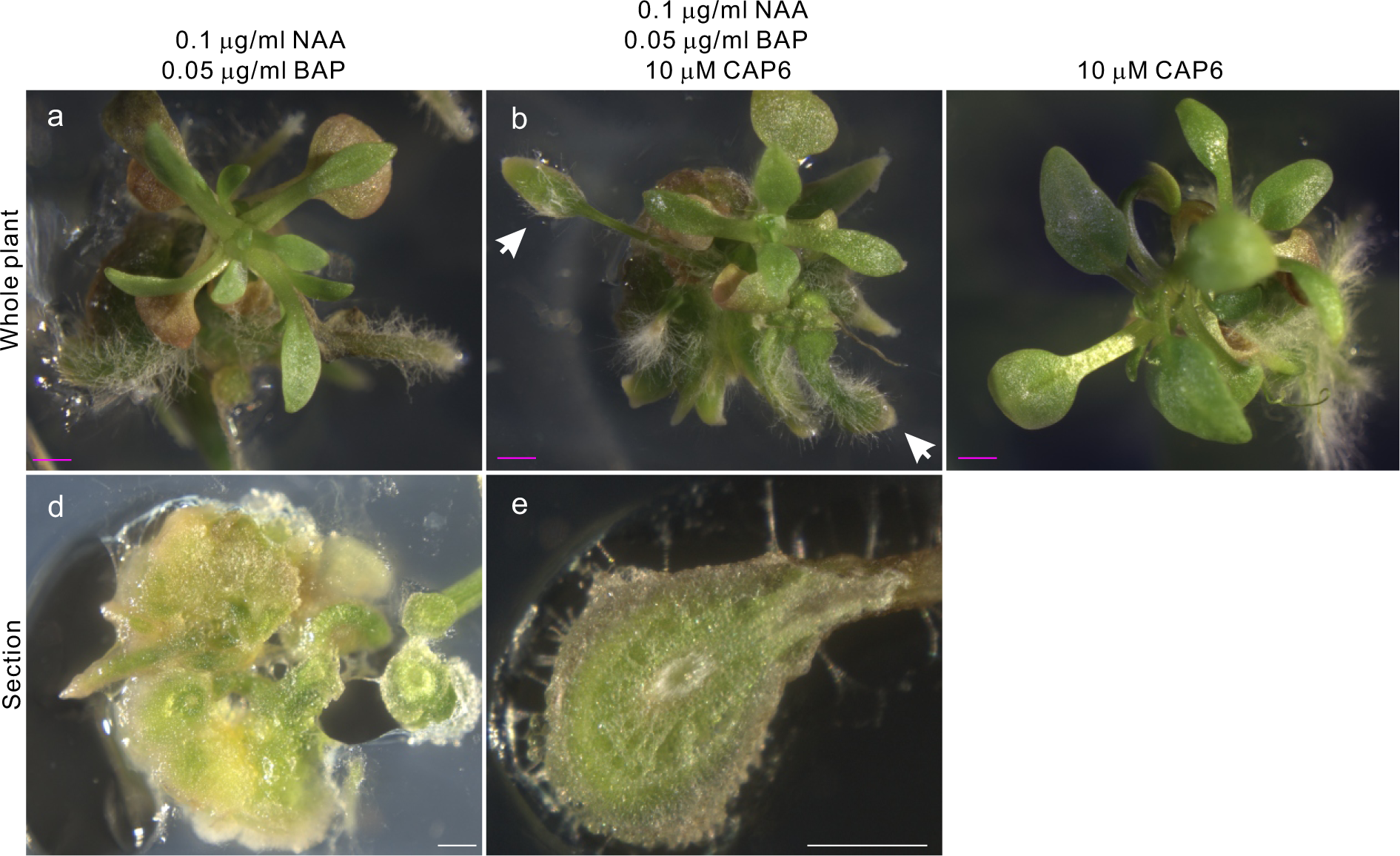
*V. peregrina* cultured on the medium with a mixture of phytohormones (0.1 μg/mL NAA and 0.05 μg/mL BAP) **(a)**, a mixture of phytohormones (0.1 μg/mL NAA and 0.05 μg/mL BAP) and 10 μM CAP6 peptide **(b)**, and 10 μM CAP-p6 peptide **(c). (b)** Arrows show artificial galls. The sections of the callus **(d)** and artificial gall **(e)** formed with a mixture of phytohormones (0.1 μg/mL NAA and 0.05 μg/mL BAP) and 10 μM CAP-p6 peptide. Bars = 1 mm (**a–c**) and 100 μm (**d** and **e**).

